# Sharing genetic admixture and diversity of public biomedical datasets

**DOI:** 10.1101/210716

**Authors:** Olivier Harismendy, Jihoon Kim, Xiaojun Xu, Lucila Ohno-Machado

## Abstract

Genetic ancestry and admixture are critical co-factors to study phenotype-genotype associations using cohorts of human subjects. Most publically available molecular datasets – genomes, exomes or transcriptomes - are however missing this information or only share self-reported ancestry. This represents a limitation to identify and re-purpose datasets to investigate the contribution of race and ethnicity to diseases and traits. we propose an analytical framework to enrich the meta-data from publically available cohorts with admixture information and a resulting diversity score at continental resolution, calculated directly from the data. We illustrate the utility and versatility of the framework using The Cancer Genome Atlas datasets indexed and searched through the DataMed Data Discovery Index. Data repositories or data contributors can use this framework to provide, as metadata, admixture for controlled access datasets, minimizing the work involved in requesting a dataset that may ultimately prove inadequate for a researcher’s purpose. With the increasingly global scale of human genetics research, research on disease risk and susceptibility would benefit greatly from the adequate estimation and sharing of admixture data following a framework such as the one presented.

## Introduction

In order to facilitate the identification and reuse of publicly available biomedical datasets, we have developed the DataMed, a search engine for data^1^. A large number of the datasets indexed and retrievable in DataMed are derived from human specimens (blood, cell lines, tissues) and contain broad genetic information (genotypes, exome, genome or transcript sequences). Using established analysis frameworks, one can extract from the raw data useful meta-data that is not necessarily collected or known from the investigators. These can include race, ancestry admixture, HLA haplotypes, telomere length, tumor viral load or purity, or cell-line identity. We present here the framework we established to efficiently call admixture on DataMed-indexed cohorts and propose to summarize and index the results through a diversity score.

Race and ethnicity have a significant influence on health and disease etiology. Whether the associated risks are due to socioeconomic, environmental or genetic factors varies among diseases and, in most of them, the associations remain to be determined. When accounting for race and ethnicity, studies generally rely on self-reporting. Self-reporting lacks accuracy to distinguish East Asian from South Asian^2^, or when subjects have strong admixture (i.e., they have 2 or more ancestries).^3,4^ The 1000 genome project has identified variants in 26 reference populations that can be grouped into 5 continental super populations^5^. From this reference dataset, one can estimate admixture in any given individual using genotypes genome-wide, or at selected Ancestry Informative Markers (AIM)^6,7^. Admixture can then be used as a covariate in genetic studies, in order to account for population structure^8^, or to identify ancestry specific signals^9^.

The availability of a uniform, genetically-based ancestry estimation for all eligible human datasets indexed in DataMed would increase their usability, allowing the selection of diverse cohorts, preparing ancestry specific meta-analysis, or simply monitoring diversity. The diversity score can facilitate the identification and assembly of ancestry specific cohorts, and enable the monitoring of racial and ethnical diversity in biomedical research datasets.

## Methods

### Data

We selected The Cancer Genome Atlas (TCGA)^10^ cohort to implement the diversity score into DataMed. Indeed this cohort is large (N=10,878), one of the most accessed cohorts in dbGAP and contains self-reported race and ethnicity. In addition, the cohort can be split into 33 sub-cohorts corresponding to each cancer type, providing an opportunity to contrast the various collections. Finally, the vast majority of samples have multiple data types (genotypes, exomes, transcriptomes), on which we can compare admixture estimation.

A total of 10,878 TCGA subjects (individuals) have been genotyped at ~10^6^ SNPs. We called admixture from the 5 continental reference populations: European (EUR), African (AFR), East Asian (EAS), South Asian (SAS) and Native American (AMR).

### Data Access and Pre-processing

The data specified below were retrieved through the National Cancer Institute (NCI) Genomic Data Commons (GDC) using the gdc-client API. We obtained the <u>*genotyping array*</u> data in the birdseed format (the result of genotype calling by birdSuite^11^), which were then converted to the plink^12^ format (MAP and PED flat files). To ensure the proper alleles were reported during the conversion, we established a relational database to decode the numeric genotype into alleles using information from the Affymetrix SNP Array 6.0 probe design and the corresponding dbSNP (v150) rsid. The <u>*RNA-Seq reads*</u> (BAM files) from the patient tumors were used to call variants using the following steps: (1) duplicate reads removal (PICARD Markduplicates), (2) split intron spanning reads (GATK v3.8), and (3) variant calling (GATK v3.8 HaplotypeCaller). We called variants from the <u>*whole exome*</u> sequence (BAM files) from the blood using freebayes^13^ (v1.1.0). For both RNA-Seq and Exome Sequencing analysis, we restricted the variant calling to known SNP (dbSNP v150) located in the exons and CDS regions of Gencode-v25^14^ respectively. The variants were filtered (DP>10 and GQ>15), and then converted to plink format using vcftools.

### Admixture analysis

For each individual, the admixture fraction for the reference population was estimated using iAdmix tool^7^. The input data were individual genotypes (MAP and PED flat files in PLINK format), and the allele frequencies from the 1000 Genomes reference populations^5^. The 1000 genome reference VCF file was based on the GRCh37 human genome build and contained allelic fractions calculated from 2,504 individuals divided into 5 super-populations: European (EUR), African (AFR), East Asian (EAS), South Asian (SAS) and American (AMR). To accommodate genotypes from different versions of the human genome reference, the SNP coordinates were converted to GRCh38 using liftOver (https://genome-store.ucsc.edu/). The output of iAdmix was a list of five admixture fractions, each with values ranging between 0 and 1. These estimates correspond to maximum likelihood estimations (MLE) through Broyden-Fletcher-Goldfarb and Shanno (BFGS), a widely used, quasi-Newton optimization method. The cumulative admixture fraction was calculated as the overall fraction of the 5 ancestries after summing up individual admixture faction across a given set of individuals. To calculate the diversity score of each cancer specific cohort, we calculated the cumulative fraction of each ancestry across all individuals in the cohort. We then computed the normalized entropy from the resulting 5 dimensional vector using R package *entropy*, as the empirical entropy divided by the maximal entropy for 5 dimensions. The benchmarking study comparing admixture determination using genotyping vs. exome vs. transcriptome was conducted on 100 subjects specifically selected in order to have a sample that maximized diversity in self-reported race and ethnicity (Supplementary Table 1).

## Results

### Admixture in TCGA data

The dominant ancestry - representing more than 80% admixture - of each individual matches well the self-reported one: 76% White Non-Hispanic are EUR dominant, 82% of Black are AFR dominant and 89% of Asian are either SAS or EAS dominant. Similarly, 53 % of subjects reported as Hispanic or Latino have at least 20% of AMR ancestry. We then determined the Cumulative Admixture Fraction (CAF) for each cancer-specific cohort (Method). The CAF reflects, at the cohort level, the fraction of total DNA from a given ancestry, rather than the fraction of individuals of a given race or ethnicity. While all cancer cohorts are predominantly EUR (Figure 1A - 46% to 93%), the fraction of non-EUR ancestry varies: Renal Cell Carcinoma (KIRP) is the cohort with highest AFR ancestry (21%), while Hepatocellular Carcinoma (LIHC) has the highest EAS ancestry (41%). While these differences may reflect the epidemiology of the disease, it is important to note that the TCGA cohort had significant ascertainment bias, including enrollment sites, tumors sizes, purity and availability requirements. Finally, in order to summarize the overall diversity of each cohort, we used the CAF to compute a normalized diversity score (DS): 0 for one ancestry only, 1 for an even fraction of all five ancestry populations. The TCGA cohorts can be ranked by decreasing diversity, revealing that Hepatocellular Carcinoma dataset as the most diverse (DS=0.7) and Uveal Melanoma as the least diverse (DS=0.22, Figure 1A). Both the diversity score and the minimal admixture level of a given ancestry can be used to filter cohorts in the DataMed index.

**Figure 1:**
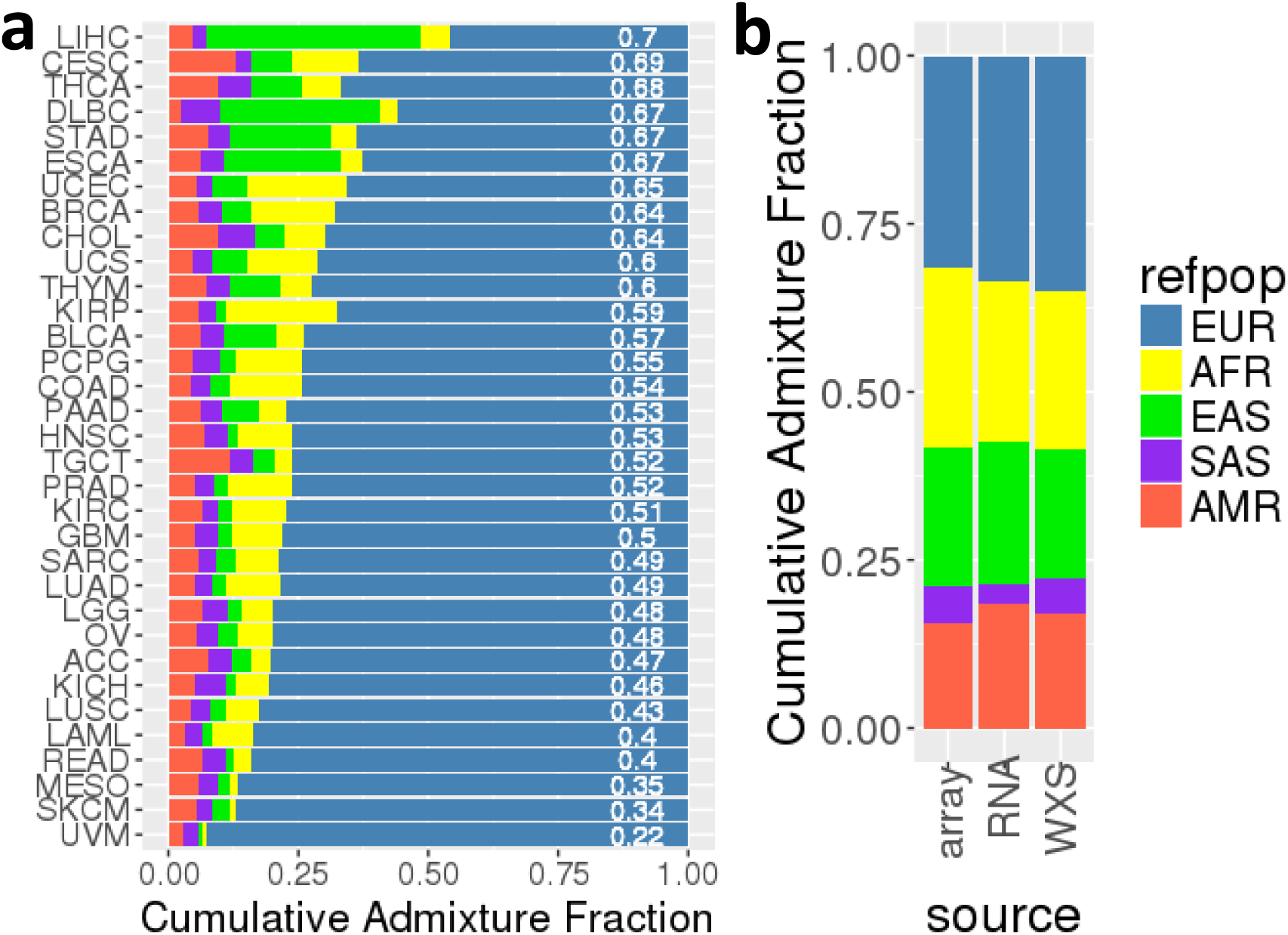
Cumulative Admixture in The Cancer Genome Atlas. **(A)** Cumulative Admixture Fraction of 33 cancer specific cohorts, inferred from the 5 reference super-populations. The cohorts are ranked by decreasing diversity score (white label) **(B)** Cumulative Admixture Fraction of a selected set of 100 diverse TCGA subjects using genotypes from genotyping array, transcriptome (RNA) or exome (WXS). <u>*Cancer Type abbreviations:*</u> Acute Myeloid Leukemia (LAML), Adrenocortical carcinoma (ACC), Bladder Urothelial Carcinoma (BLCA), Brain Lower Grade Glioma (LGG), Breast invasive carcinoma (BRCA), Cervical squamous cell carcinoma and endocervical adenocarcinoma (CESC), Cholangiocarcinoma (CHOL), Colon adenocarcinoma (COAD), Esophageal carcinoma (ESCA), Glioblastoma multiforme (GBM), Head and Neck squamous cell carcinoma (HNSC), Kidney Chromophobe (KICH), Kidney renal clear cell carcinoma (KIRC), Kidney renal papillary cell carcinoma (KIRP), Liver hepatocellular carcinoma (LIHC), Lung adenocarcinoma (LUAD), Lung squamous cell carcinoma (LUSC), Lymphoid Neoplasm Diffuse Large B-cell Lymphoma (DLBC), Mesothelioma (MESO), Ovarian serous cystadenocarcinoma (OV), Pancreatic adenocarcinoma (PAAD), Pheochromocytoma and Paraganglioma (PCPG), Prostate adenocarcinoma (PRAD), Rectum adenocarcinoma (READ), Sarcoma (SARC), Skin Cutaneous Melanoma (SKCM), Stomach adenocarcinoma (STAD), Testicular Germ Cell Tumors (TGCT), Thymoma (THYM), Thyroid carcinoma (THCA), Uterine Carcinosarcoma (UCS), Uterine Corpus Endometrial Carcinoma (UCEC), Uveal Melanoma (UVM). <u>*Super-Populations abbreviations*</u>: European (EUR), African (AFR), East Asian (EAS), South Asian (SAS), Native American (AMR).

### Assessing admixture using transcriptomes or exomes

A large number of studies indexed by DataMed may not contain readily available genotype information. This is particularly the case for studies generating whole exome or whole transcriptome. In order to expand the utility of our approach to these cohorts, we evaluated the admixture and diversity score estimation using also exome and transcriptome data and compared them to results from the genotyping array. For this comparison we selected 100 TCGA subjects representing all possible self-reported race and ethnicities to ensure the results would be consistent across various genetic backgrounds (Supplementary Table 1). After variant calling and filtering (Methods), we identified a median of 21,327 and 838 usable variants in the exome and transcriptome of each subject, respectively. The ancestries with maximum admixture were consistent across for all three methods for 82/100 subjects. The subjects with inconsistent results were more admixed based on the genotyping array results (maximum admixture 0.89±016 vs. 0.72±022). As a result, the CAF estimated from the exome or transcriptome variants were consistent with the ones from genotyping array (r=0.97 for both Figure 1B) and all three diversity scores were similar - 0.93, 0.92, 0.90 for genotyping, exome and transcriptome, respectively.

## Discussion

A number of studies agree that the genetic ancestry is far more accurate and therefore superior to self-reported race and ethnicity^15–17^. To date, it is not possible to get an accurate estimation of the racial and ethnical diversity of a cohort before looking inside the dataset (i.e., looking at the individual level data) and calling admixture. A low resolution, 5 super-populations admixture estimate like the one we present here is very valuable for investigators who want to account for admixture in their genetic studies or select patients to assemble a cohort for meta-analysis of a given ancestry. In order to calculate the diversity score, we had to request access to the cohort for this specific task, a step that may not be permitted for certain cohorts or that is not necessarily scalable. However, the diversity score does not have to be generated by the DataMed team, but could instead be computed by the data owners and shared as an additional piece of metadata that could be used downstream for cohort selection.

The admixture and diversity score generated are well applicable on a variety of broad molecular datasets. We demonstrated their validity from exome and transcriptome. To date, 176×10^3^ and 201×10^3^ human transcriptome (RNA-Seq) and exome datasets, respectively, are hosted by the NCBI Sequence Read Archive (SRA). Among those, 82% of transcriptomes and 12% of exomes are available without restriction, and likely none of them have associated genetic ancestry information. Beyond transcriptome or exomes, ancestry can also be called from ChIP-seq datasets from human sample - more than 31×10^3^ currently available in the NCBI SRA. A typical ChIP-Seq dataset may cover 10^6^ bp genome, harboring 1000 SNPs, the majority of which have been genotyped in the 1000 genome reference populations, representing a sufficient number to determine genetic admixture.

The same way the Gene Expression Omnibus has the ability to search and rank datasets based on differential expression of a specific gene, one can hope that future, innovative data sharing strategies will include as many of such data-derived features, like genetic admixture, generated in an automated, standardized way at the time of the deposition. The relative simplicity of calling admixture on molecular datasets may encourage more careful analytical design. While we know that germline genetics may play a role in disease etiology or phenotypic differences, it is rarely taken into account in pre-clinical or clinical studies. Using admixture from known continental ancestry as a first-order surrogate for germline genetic differences, one could account for this important co-variate and relate it to a population trait. In the past, pre-clinical studies based on a small number of cell lines or samples could not reasonably account for inherited genetic variation. Nowadays, pre-clinical studies are becoming larger and more systematic, such as the Cancer Cell Line Encyclopedia^18^ (N=750 cell lines), but to our knowledge they still do not account for genetic ancestry. More recently, genetically diverse sets of lymphoblastoid cell lines^19^ or induced pluripotent stem cells^20^ have been made available for research, documenting the increasing interest in performing pre-clinical research in large sets of genetically diverse samples and cell lines. The availability of genetic admixture as a piece of metadata in the public datasets would therefore increase their utility to design analysis accounting for inherited genetic background, resulting in more accurate statistical models and in a better understanding of the contribution of genetic variation and ancestry to disease etiology or drug response. The diversity score featured in the DataMed index provides an optimal way for researchers to select the adequate datasets for this task, without the need to disclose individual level data.

## Acknowledgements

This work was supported by a grant from NIH/NIAID (U24AI117966) to L.O.-M.

## References

1. Ohno-Machado, L. et al. Finding useful data across multiple biomedical data repositories using DataMed. Nat. Genet. 49, 816–819 (2017).

2. Mersha, T. B. & Abebe, T. Self-reported race/ethnicity in the age of genomic research: its potential impact on understanding health disparities. Hum. Genomics 9, 1 (2015).

3. Rishishwar, L. et al. Ancestry, admixture and fitness in Colombian genomes. Sci. Rep. 5, 12376 (2015).

4. Sucheston, L. E. et al. Genetic ancestry, self-reported race and ethnicity in African Americans and European Americans in the PCaP cohort. PLoS One 7, e30950 (2012).

5. Auton, A. et al. A global reference for human genetic variation. Nature 526, 68–74 (2015).

6. Alexander, D. H., Novembre, J. & Lange, K. Fast model-based estimation of ancestry in unrelated individuals. Genome Res. 19, 1655–1664 (2009).

7. Bansal, V. et al. Fast individual ancestry inference from DNA sequence data leveraging allele frequencies for multiple populations. BMC Bioinformatics 16, 4 (2015).

8. Halder, I. & Shriver, M. D. Measuring and using admixture to study the genetics of complex diseases. Hum Genomics 1, 52–62 (2003).

9. Fejerman, L. et al. Admixture mapping identifies a locus on 6q25 associated with breast cancer risk in US Latinas. Hum. Mol. Genet. 21, 1907–1917 (2012).

10. Network, T. C. G. A. R. et al. The Cancer Genome Atlas Pan-Cancer analysis project. Nat Genet 45, 1113–1120 (2013).

11. Korn, J. M. et al. Integrated genotype calling and association analysis of SNPs, common copy number polymorphisms and rare CNVs. Nat Genet (2008). doi:ng.237 [pii]10.1038/ng.237 [doi]

12. Purcell, S. et al. PLINK: a tool set for whole-genome association and population-based linkage analyses. Am J Hum Genet 81, 559–575 (2007).

13. Garrison, E. & Marth, G. Haplotype-based variant detection from short-read sequencing. 9 (2012).

14. Harrow, J. et al. GENCODE: The reference human genome annotation for The ENCODE Project. Genome Res. 22, 1760–1774 (2012).

15. Spector, S. A. et al. Genetically determined ancestry is more informative than self-reported race in HIV-infected and -exposed children. Medicine (Baltimore). 95, e4733 (2016).

16. Smith, E. N. et al. Genetic ancestry of participants in the National Children’s Study. Genome Biol. 15, R22 (2014).

17. Lee, Y. L., Teitelbaum, S., Wolff, M. S., Wetmur, J. G. & Chen, J. Comparing genetic ancestry and self-reported race/ethnicity in a multiethnic population in New York City. J. Genet. 89, 417–423 (2010).

18. Barretina, J. et al. The Cancer Cell Line Encyclopedia enables predictive modelling of anticancer drug sensitivity. Nature 483, 307–603 (2012).

19. Altshuler et al., D. & Consortium, T. I. H. A haplotype map of the human genome. Nature 437, 1299–1320 (2005).

20. Panopoulos, A. D. et al. iPSCORE: A Resource of 222 iPSC Lines Enabling Functional Characterization of Genetic Variation across a Variety of Cell Types. Stem cell reports 8, 1086–1100 (2017).

